# Revisiting Analog Electrical Stimulation with Current Focusing in a Guinea Pig Model of Cochlear Implants

**DOI:** 10.64898/2026.07.02.735566

**Authors:** V. Adenis, RA. Bartholomew, JI. Lee, A. Jung, MC. Brown, DJ. Lee, JG. Arenberg

## Abstract

Modern cochlear implants (CIs) use pulsatile stimulation to restore hearing for individuals with severe hearing loss. CIs provide robust speech recognition in quiet but poorly represent temporal fine structure (TFS), needed for challenging listening situations. Analog stimulation preserves the acoustic waveform and may better encode TFS, yet it has not been evaluated combined with modern current-focusing strategies. We compared neural responses in the inferior colliculus (IC) evoked by CI stimuli consisting of 100 pulses/s biphasic pulse trains and 100 cycles/s sinusoidal analog stimulation with monopolar, bipolar, and tripolar electrode configurations in urethane-anesthetized guinea pigs. Following cochlear implantation, multiunit activity was recorded from the tonotopic axis of the central nucleus of the IC using 16-channel silicon probes. Detection thresholds, spread of excitation, vector strength, sustained response percentage, and temporal response properties were quantified. Analog stimulation consistently evoked significantly lower activation thresholds than pulsatile stimulation while maintaining comparable or sometimes narrower spatial selectivity across stimulation modes. In contrast, analog stimulation generated lower vector strength, larger tonic response components, and a pronounced level-dependent polarity effect. At low stimulus levels, responses were dominated by the cathodic phase of the sinusoidal waveform, whereas increasing stimulus level responses were elicited by both phases, producing synchronization at twice the stimulus frequency. These findings demonstrate that stimulation waveform strongly influences temporal coding while having relatively little effect on the spatial distribution of neural activation. These results provide a physiological basis for reexamining analog stimulation as an alternative strategy for cochlear implant sound coding.

## Introduction

Cochlear implants (CIs) are widely regarded as one of the most successful neural prostheses. Average CI users achieve 70–80% accuracy in open-set sentence recognition in quiet environments (1), but challenges persist regarding understanding speech in noise, pitch perception (2,3), and music appreciation (4,5). Hearing restoration through CIs relies on two core aspects: how the acoustic signal is translated into electrical patterns (coding strategy), and how those patterns are spatially distributed within the cochlea through the electrode array (electrode configuration).

CIs process sound frequency through a series of bandpass filters that selectively deliver stimulation to the multiple, tonotopically arranged electrodes. The most common type of sound coding strategy used in modern CIs employs pulsatile stimulation and some version of the Continuous Interleaved Sampling (CIS) (6) strategy that temporally interleaves stimulation pulses across electrodes so that only a single electrode is active at a time, thereby preventing electrical field summation and likely reducing channel interaction. This pulsatile strategy has proven highly effective for speech perception in quiet; however, it provides only coarse (if any) representations of intricate sound features such as temporal fine structure (TFS), which limits performance in noisy environments or for complex tasks. In normal acoustic hearing, TFS cues are preserved for frequencies up to approximately 1.5kHz, and are represented by the synchronized spiking of auditory nerve fibers in response to pure tones or amplitude-modulated tones (7). In CI hearing with the CIS-based strategy, TSF cues are not preserved because of the use of a fixed carrier rate (of at least 500 pulses/s (pps)). Only slow changes in envelope amplitude are available. Over the last decade, alternative sound coding strategies have been developed to represent some aspects of TFS, but their efficacy for complex stimulus perception has not been demonstrated (8,9) and the lack of TFS in other available sound coding strategies remains a fundamental limitation of CI outcomes (10–12).

An alternative sound coding strategy based on Analog Stimulation (AS), also called Simultaneous Analog Stimulation or SAS (13,14), more faithfully represents the acoustic waveform by delivering a continuous electrical signal (15). In this strategy the envelope and all aspects of the TFS are encoded directly in the stimulus waveform, preserving the instantaneous amplitude variations of the filtered signal by translating them directly into a time-varying electrical waveform. This theoretically enables the auditory nerve to respond to the stimulus in a manner analogous to normal hearing.

For a time, a subset of CI systems offered both CIS and SAS to recipients, allowing for direct clinical comparisons between the two strategies. A plethora of early psychoacoustic studies found that SAS yielded overall compared speech perception performance similar to CIS alone (as summarized in Stupak et al. 2018 (16)) and a meaningful subset of users performed better with, and preferred, the analog stimulation strategy. For example, Koch et al. reported that one third of patients preferred to use the SAS strategy after three months of implant experience; although mean sentence scores were similar between groups, word recognition scores were significantly higher for SAS users than for CIS users (17). Similarly, another study found that half of subjects selected SAS as their preferred strategy, with the use of SAS resulting in higher overall patient performance (18).

A promising but largely unexplored synergy would be the combination of AS with modern current focusing techniques such as bipolar and tripolar mode (19,20). These techniques use adjacent intracochlear electrodes in the linear array of the cochlear implant to spatially restrict the electrical field and likely reduce channel interactions. If the primary historic shortcoming of AS was the summation of simultaneous electrical fields across adjacent channels, then appropriately calibrated bipolar or tripolar configurations offer a principled means of reducing that interaction while preserving the TFS that gives AS its inherent advantage over pulsatile strategies. The tripolar configuration effectively reduces channel interaction (21), enabling the transmission of temporal information by reducing distortion from the activation of overlapping population of neurons by multiple channels (22). In early AS some clinicians successfully fitted patients with the SAS strategy using the bipolar mode (18), and CI listeners who preferred SAS under this configuration were found to have electrode-to-modiolus distances that were smaller, suggesting that proximity and, likely, more restricted electrical fields may be key moderators of AS success. A recent group attempted to overcome the shortcomings of conventional strategies by utilizing within-channel encoding of temporal cues through novel analog, single channel encoding strategies and computational modeling (23). The present study evaluates the coupling of modern current focusing strategies with AS in a guinea pig model to assess its efficacy as a TSF-preserving alternative to pulsatile interleaved strategies.

## Methods

### Animals

Eleven Hartley guinea pigs (*Cavia Porcellus*) were used in this study. At the time of surgery, age ranged from 4 to 10 months, weight from 350 to 850 g and both sexes were used. Procurement, housing and all experimental procedures were performed in accordance with the National Institutes of Health guidelines for the care and use of laboratory animals as well as the approved animal care and use protocols at Mass Eye and Ear, Boston, MA. All animals used in this study had normal hearing confirmed with neurophysiological responses (see below (19,24,25)).

### Surgical approaches

Surgeries were performed under general anesthesia induced by an initial dose of urethane (900 mg/kg, i.p.) and fentanyl (0.15 mg/kg, i.p.) and supplemented by partial doses when reflex movements were observed after pinching the hind paw. Each animal also received a one-time dose of Haloperidol (7.5-10 mg/kg, i.p.) and atropine (0.04 mg/kg, i.p.) at induction. A heating pad (Harvard Apparatus™) maintained body temperature at approximately 37 °C. Vitals (heart rate, SpO2, body temperature) were monitored throughout the experiment every 30-60 minutes. After anesthesia induction, animals were placed in a stereotaxic frame (Kopf) and received an injection of local anesthetic (Lidocaine 1%, s.c.). Skin was incised, and the temporal muscles pushed aside such that the skull could be cleaned and dried. Three stainless steel screws were threaded into the skull via burr holes to anchor a miniature headplate. Screws and headplate were then embedded in acrylic dental cement to fix the animal’s head. All the following procedures were conducted with a binocular surgical microscope (Zeiss).

A craniotomy was performed on the right side of the skull, above the visual cortex. After removing dura, the overlying cortex was aspirated to expose the underlying Inferior Colliculus (IC). A 16-channel recording probe (A16 single shank, 100 µm spacing, NeuroNexus Technologies, Michigan) was then inserted ∼3 mm deep into the IC to target the central nucleus (Fig. 1A). The silicon recording probe was inserted along the dorso-ventral axis, with a 30° angle in the coronal plane to sample neuronal activity along the tonotopic axis of the IC. Placement was confirmed by recording pure tone Frequency Response Areas (FRAs, Fig. 1A, see below). Placement of the IC probe was adjusted accordingly until frequency tunning curves could be obtained at each recording site from a range of 8 to 45 kHz best frequencies. The recording probe was then fixed by filling the cavity with 10% agarose gel and a layer of acrylic dental cement connected to the headplate. After fixing the recording probe, final, more detailed FRAs were measured.

**Figure 1:**
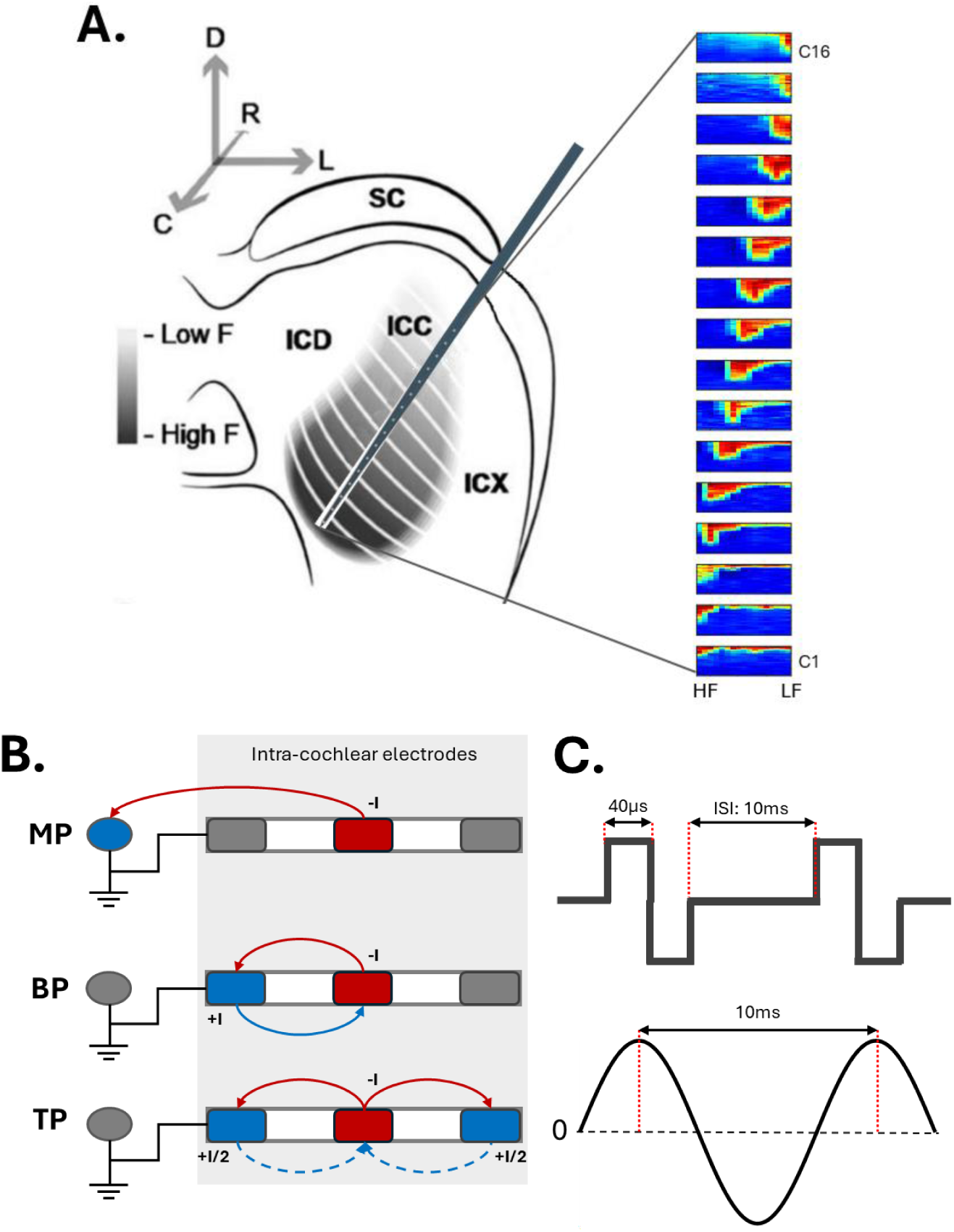
A Left: Representation of the 16-channel recording probe insertion along the tonotopic axis of the guinea pig inferior colliculus (coronal view). Derived from Lenarz et al. 2006, SC; superior colliculus, ICD: dorsal inferior colliculus, ICC: central inferior colliculus, ICX: inferior colliculus external cortex. Drawing and probe not to scale. Right: Examples of frequency response areas recorded under the 16-channel recording probe when presenting pure tones from 8 to 45 kHz (abscissa) from 0 to 90 dB SPL (ordinate). Color code represents the normalized firing rate (red: high rate; blue: low rate). B. Depiction of the three stimulation modes used in the study. MP: monopolar, one intracochlear electrode (red) delivers the current while the external ball electrode serves as the return electrode (blue). BP: bipolar, two neighboring intracochlear electrodes serve as active (red) and return (blue) electrodes while delivering mirrored stimuli of equal level. TP: tripolar, a central electrode delivers current (red) while surrounding electrodes on each side (blue) deliver mirrored stimuli, each at half the current level of the central one. C. Top: Pulsatile stimulation consisted of symmetric biphasic square wave pulses (40 us pulse width with an inter-stimulus interval of 10 ms). Bottom: Analog stimulation consisted of charge-balanced sinusoidal stimulation (10 ms period).

The left cochlea was exposed with a postauricular incision followed by blunt dissection of overlying muscles to expose the otic bulla. The bulla was opened with a surgical drill until the cochlea round window (RW) was exposed. To lesion hair cells and avoid unwanted outer hair cell activity induced by electrical stimulation (e.g. (26,27)), two 10% Neomycin infusions (10 mg/ml) were made through the RW, 20 minutes apart (total action time: 40 to 45 min). Briefly, a micro syringe was inserted in the cochlea and the Neomycin injected until egress was observed. An 8-electrode cochlear implant (Cochlear Ltd., Platinum-Iridium, half-banded contact diameter: 400µm, inter-electrode distance: 500 µm center-to-center) was then inserted in the scala tympani through the RW until resistance was met. In most cases, the eighth electrode was inside the RW but still visible. An extra-cochlear ground was placed in the neck muscles and the implant head-connector secured on the headplate with cyanoacrylate adhesive (Vetbond, 3M, Minnesota).

### Frequency Response Areas (FRAs)

Stimulus generation, delivery, and neural recordings were under the control of a PXI computer (National Instruments) and custom Labview/Matlab interfaces developed at Mass Eye and Ear (system characteristics and codes can be shared upon request and can be viewed on Mass. Eye and Ear public GitHub)

FRA were constructed by presenting 30 ms duration tone bursts from 8 to 45 kHz, in quarter octave steps, at a rate of 4 burst/s. Stimulus levels ranged from 0 to 90 dB SPL in 5dB steps. Each stimulus was repeated 8 times, and the associated responses averaged to create the FRAs for each of the 16-recording channel (see below for signal extraction).

### Electric stimulation

This study reports the evoked responses obtained with pulsatile and analog stimulation combined with current focusing strategies: mono-, bi- and tripolar pulse trains (MP-P, BP-P and TP-P) and their equivalents with AS (MP-A, BP-A, and TP-A). All stimuli were charge-balanced to avoid ionic imbalance that could damage spiral ganglion neurons. All three configurations of stimulation described below have been widely used in the literature (e.g. (28,29)). Specifically, the monopolar configuration uses one intra-cochlear active electrode and the extra-cochlear electrode as return electrode. Bipolar and tripolar also use one intra-cochlear active electrode but use respectively one or two neighboring electrodes for returning current. For bipolar, return was always the basal neighbor while tripolar used the two flanking neighbors (one apical and one basal) as returns, and shared current equally (Fig. 1B).

Two paradigms of stimulation were used: pulsatile and analog (Fig. 1C). Pulses were cathodic-first, 40 μs/phase duration with no interphase gap, and presented in 30 ms duration burst at a rate of 100 pulses/s (pps). Each pulse train had an Inter-stimulus Interval (ISI) of 1 s and levels of stimulation ranged from -46 to 6 dB re 1V (dB V). Analog stimulation was continuous, cathodic-first sinusoidal stimulation, 30 ms duration, with an ISI of 1 s (Fig. 1C), equivalent to 100 cycles/s (cps). A subgroup of animals was also stimulated with analog stimulation alternating in polarities with cathodic and anodic phase first presented every odd and even repetition, respectively. Level of stimulation ranged from -46 to 6 dB re 1V (dB V).

### Inferior Colliculus Multi-Unit Activity (MUA) recordings and analysis

MUA were recorded in a similar fashion as in McInturff et al. 2023, 2022 (30,31). Briefly, raw voltage signals from the IC recording channels were sampled at 25 kHz. An average of activity across all 16 recording channels was used as a global reference and subtracted from each single channel waveform to remove large artifacts present across channels (e.g. heartbeat, breathing). Waveforms were then stimulus-aligned. For pulsatile stimulation, stimulus artifacts were also minimized by zeroing the first 0.7 ms following each pulse. For each channel, the average of the aligned waveform was then used as a local reference and subtracted from each individual waveform to remove local field potential (LFP) components still present in the signal. The resulting waveforms were then digitally filtered (5th order, Butterworth bandpass filter, 300 Hz to 3 kHz).

Multiunit response latencies were determined as the timestamps of local maxima above a voltage threshold. A 0.8 ms dead time was implemented following each event (32) to avoid counting events that are not biologic. Voltage thresholds for spike triggering were calculated separately for each channel as 4x the standard deviation of voltage traces during the stimulus-off period (0.5 s). Post-stimulus time histograms (PSTHs) with bin widths of 0.6 ms to appropriately capture the sub-millisecond temporal precision of IC neurons (33,34), were constructed to study MUA offline, post-experiment.

### Data analysis

#### D-prime

Offline analyses started with the characterization of thresholds and saturation levels, and “stimulation tuning curves” (STCs). We then conducted cumulative d-prime analysis of the PSTHs (*d’*) as previously described in Arenberg, Bierer and Middlebrooks 2010 (35). Briefly, for each stimulus-level combination, a receiver-operating characteristic (ROC) curve was formed from the trial-by-trial PSTHs. For each channel, the average PSTH at a given level was considered the signal condition while the background condition consisted of the PSTH for the level below. The area under the ROC curve (AUC) yielded the proportion of trials in which the current level was discriminable from the previous level. That proportion was *z*-scored and multiplied by √2 to give the discrimination index (*d*′). Cumulative *d’* was finally defined by the summation of the current *d’* and *d’*-1 level. A log-linear correction was applied to avoid any extreme AUC value that could bias the analysis (36). Detection threshold was taken as the interpolated stimulus level at which cumulative *d*′ = 2 for each recording site.

#### Spread of Excitation

The spread of activation along the tonotopic axis of the IC was represented by stimulation tuning curves (STCs), with a response contour of *d*′ = 2 as a function of the depth along the recording probe (vertical axis) and stimulus current level (horizontal axis). This iso-rate curve was then used to calculate the Spread of Excitation (SoE) as follows. The recording channel with the lowest level to achieve *d’*>2 served as the tip of the STC, then the SoE width was calculated as the distance (channel span) between this iso-rate curve boundary at 3 dB above response threshold.

#### Vector Strength and Sustained Response Percentage

Vector strength (VS) was calculated as described by Goldberg & Brown (1969) (37) and its statistical significance assessed by the Rayleigh test (Yin & Kuwada, 1983 (38–40)) with a significance threshold of R<0.05 after applying a correction similar to Chung et al., 2014 (41)). PSTHs reported in this study contained a sustained response, tonic component in addition to the synchronous response. The sustained activity reduced the VS and distorted the analysis. To estimate and remove the tonic component, neural activations were re-distributed in time 500 times, were averaged, and subsequently subtracted from the PSTHs.

The proportion of sustained response within the PSTHs (or Sustained Response Percentage, SRP) was also quantified as follows: PSTHs were lightly smoothed with a quadratic fit (through the MATLAB function *smooth* with the ‘loess’ method) and their local peaks (maxima) and valleys (minima) identified. Valleys were averaged and reported as percentages using average maxima as the reference value.

#### Euclidian distance and angle

Within each IC recording of the 16-channel array, Euclidian Distances (EuD, in a.u.) and Angles (EuA, in degrees) were calculated for each recording channel (Ch) in relation to the local threshold as follows:

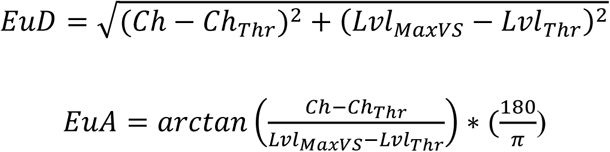

Where ChThr is the channel of the array with the lowest threshold and Ch is the channel being analyzed. LvlThr and LvlMaxVS followed the same logic with the addition that LvlMaxVS represents the level of stimulation associated with the maximum VS for this channel.

### Statistical Analysis

All statistical analyses were performed using MATLAB R2025b (MathWorks, Natick, MA, USA). Group data are presented as median ± interquartile range (IQR) given non-normal distributions (tested through Lilliefors tests). To compare pulse and analog stimulation modes, rank-based permutation tests were used (10,000 permutations, α = 0.05, two-tailed). For threshold and spread of excitation measurements (one value per recording configuration), simple rank-based permutation tests were performed. For vector strength, Euclidian distances and angles (multiple values per configuration), hierarchical permutation tests were instead used where group assignments were permuted at the animal level to account for within-animal correlations to avoid pseudo-replication. This approach is conservative and controls the non-independence of measurements from the same animal. Note that while mixed-effects models are powerful for hierarchical data, they assume specific distributions (typically normal or log-normal). Preliminary analyses revealed strong deviations from normality that could not be corrected by standard transformations. Rank-based permutation tests provide a distribution-free alternative that maintains appropriate Type I error rates while accounting for the hierarchical structure through animal-level permutation. That approach was verified as appropriately conservative by comparing results with standard Mann-Whitney tests for the subset of data where independence could be assumed. All tests were Benjamini-Hochberg corrected when appropriate.

## Results

### Analog stimulation evokes responses with narrower extent of response and lower activation thresholds

Representative stimulation tuning curves for monopolar, bipolar and tripolar (Fig. 2A) show that responses to monopolar stimulation are the broadest, those to bipolar are narrower, and those to tripolar are the narrowest. For example, the responses to monopolar pulsatile stimulation (MP-P, top left in Fig. 2A) were about the same for all recording channels, whereas the responses to tripolar pulsatile stimulation (TP-P, top right in Fig. 2A) had large responses for a narrow band of recording channels (channels 9-13). Also, in general, the responses to pulsatile stimulation (Fig. 2A) were broader than those to analog stimulation (Fig. 2B). In fact, the narrowest pattern, for TP-A stimulation, was V-shaped with a small tip region composed of responses from recording channel 10. In contrast, monopolar stimulation resulted in broad, virtually untuned response areas for both pulsatile and analog stimulation. Another difference was that thresholds were lower for analog stimulation compared to pulsatile stimulation (note different dB axis scales for Fig. 2A vs. 2B). This difference in threshold is quantified in Fig. 2C. As stimulus configuration became more focused (from mono- to tripolar analog), response thresholds increased for all stimulus types (pulse train and analog; all p-values < 0.005).

**Figure 2:**
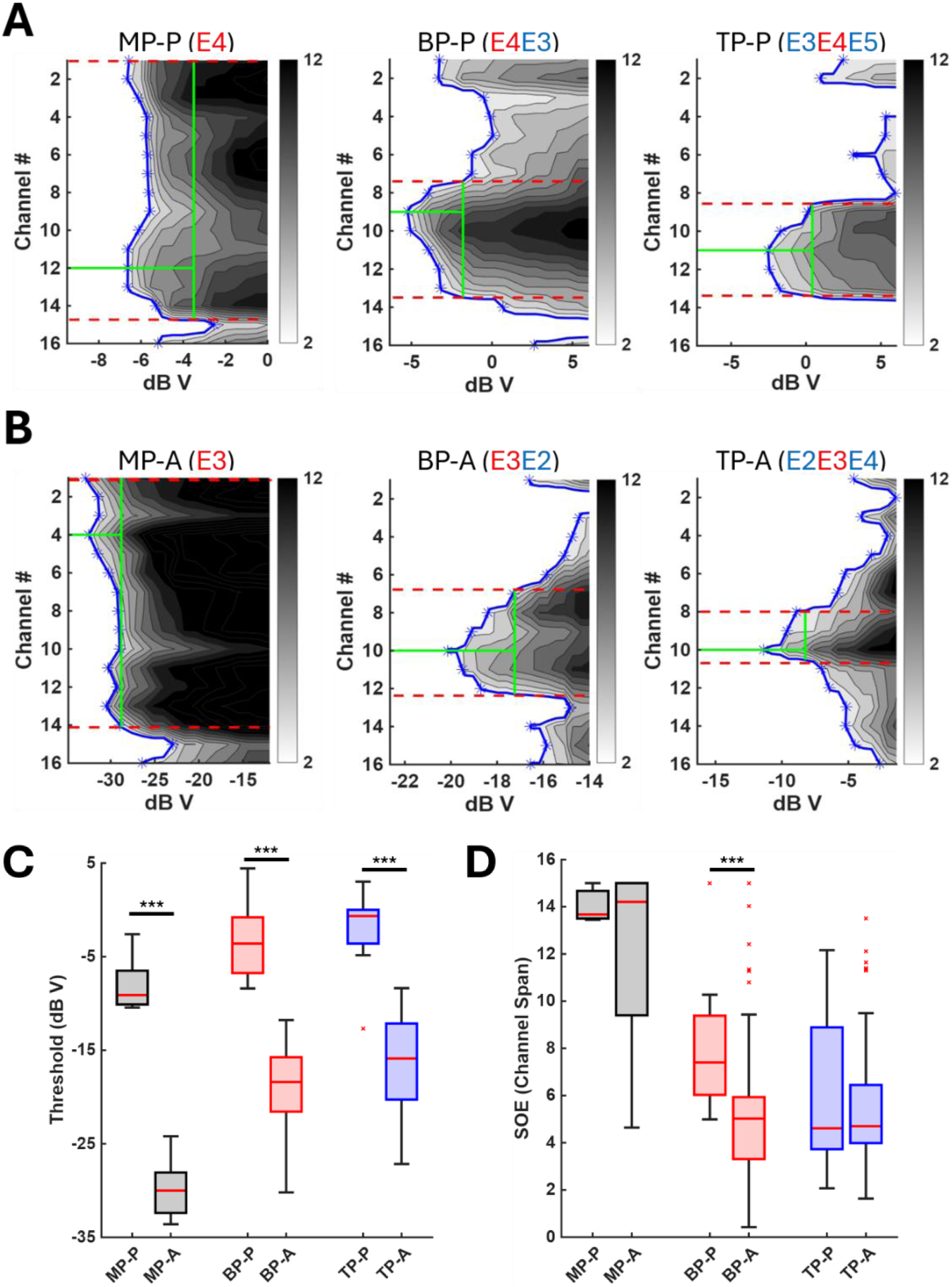
Spatial profile of IC responses. A. Pulsatile stimulation responses: D-prime isocurve maps observed across the recording array for mono-, bipo- and tripolar 100 pps pulse trains (MP-P/left, BP-P/middle and TP-P/right respectively) as a function of stimulation level while using the same active electrode. Blue curve and markers represent the threshold isocurve. Green vertical lines represent the spread of excitation measured at 3dB above threshold. Red horizontal dashed lines represent SoE borders (Left: 13.76, middle: 6.1 and right: 4.82). B. Analog stimulation responses: D-prime isocurve maps observed across the recording array for mono-, bipo- and tripolar 100 cps analog stimulation. Representations are following same conventions as in A (Left: MP-A, SoE= 13.12; middle: BP-A, SoE= 5.49; right: TP-A, SoE= 2.59, respectively) C. Distribution of evoked responses thresholds observed in the IC across all tested mode of stimulation (grey: monopolar, red: bipolar, blue: tripolar; nomenclature is identical to A). All statistical tests were rank-based permutation tests and significance is reported through p-values as follows, *p<0.05, **p<0.01, ***p<0.001. Note how analog stimulation was always linked with lower thresholds of activation, all the while following the previously described trend of more focused stimulation mode being associated with higher thresholds. D. Distribution of the Spread of Excitation recorded in the IC across all tested stimulation modes (same nomenclature as in C, significance is reported in similar fashion). Although pulsatile and analog remain equivalent, analog stimulation can evoke smaller SOEs for mono- and bipolar than pulsatile stimulation (MP-A and BP-A vs. MP-P and BP-P). Tripolar analog (TP-A) showed less variability than TP-P but achieved similar SoE on average.

To quantify the extent of response, the SoE at 3dB above threshold was calculated for each response area (green vertical lines on Fig. 2A,B). For monopolar, bipolar, and tripolar pulsatile stimulation, the examples shown (Fig. 2A) have spans of 13.76, 6.1, and 4.8. For monopolar, bipolar, and tripolar analog stimulation, the examples shown (Fig. 2B) have spans of 13.12, 5.49, and 2.59. Group data for SoE are plotted in Figure 2D. Within the same mode of stimulation, analog stimulation always had lower thresholds than pulse stimulation (all p-values < 0.0001). As stimulus configuration became more focused (from mono to tripolar analog), response areas tended to become narrower, particularly with AS (p-values < 0.0001 for MP-A vs. BP-A and MP-A vs. TP-A, p-value= 0.84 for BP-A vs. TP-A). SoE was significantly smaller for bipolar analog (BP-A) compared to its pulse equivalents (p-value < 0.0001, BP-A vs. BP-P) but tripolar analog (TP-A) and monopolar analog (MP-A) remained similar to their pulse equivalents (TP-P and MP-P) despite MP-A showing a greater variability toward smaller SoE values (all p-values < 0.05).

### Temporal profile: Responses to Analog stimuli show a strong Polarity Effect

Responses to both pulsatile and analog stimulation showed strong temporal synchrony to the stimulation (Fig. 3). However, responses to analog stimulation (Fig. 3, right) have a strong, level-dependent polarity effect (PE) that was not observed with pulse-train stimulation Fig. 3, left). This effect was seen for individual channels (Fig. 3A,B) and for average responses for all channels (Fig. 3C). At threshold levels, responses were predominantly evoked by the second, cathodic phase of the analog sinusoid (delimited by the red sections of Fig 3.A-right and in green PSTH of Fig 3.B-right), and as stimulation level increased, responses appeared for both phases (as indicated by the presence of spikes in both red and blue sections of Fig 3.A-right and by the darker PSTHs in Fig 3.B-right and C-right). Average PSTHs obtained separately with either anodic-first or cathodic-first analog stimulations, shown in Figure 3.D, revealed lower thresholds for the cathodic phase regardless of which phase was presented first. Another characteristic of the analog-evoked response of note is the presence of off-responses approximately 10 ms after the end of stimulation in about 80% of the cases. Pulse-evoked responses had none.

**Figure 3:**
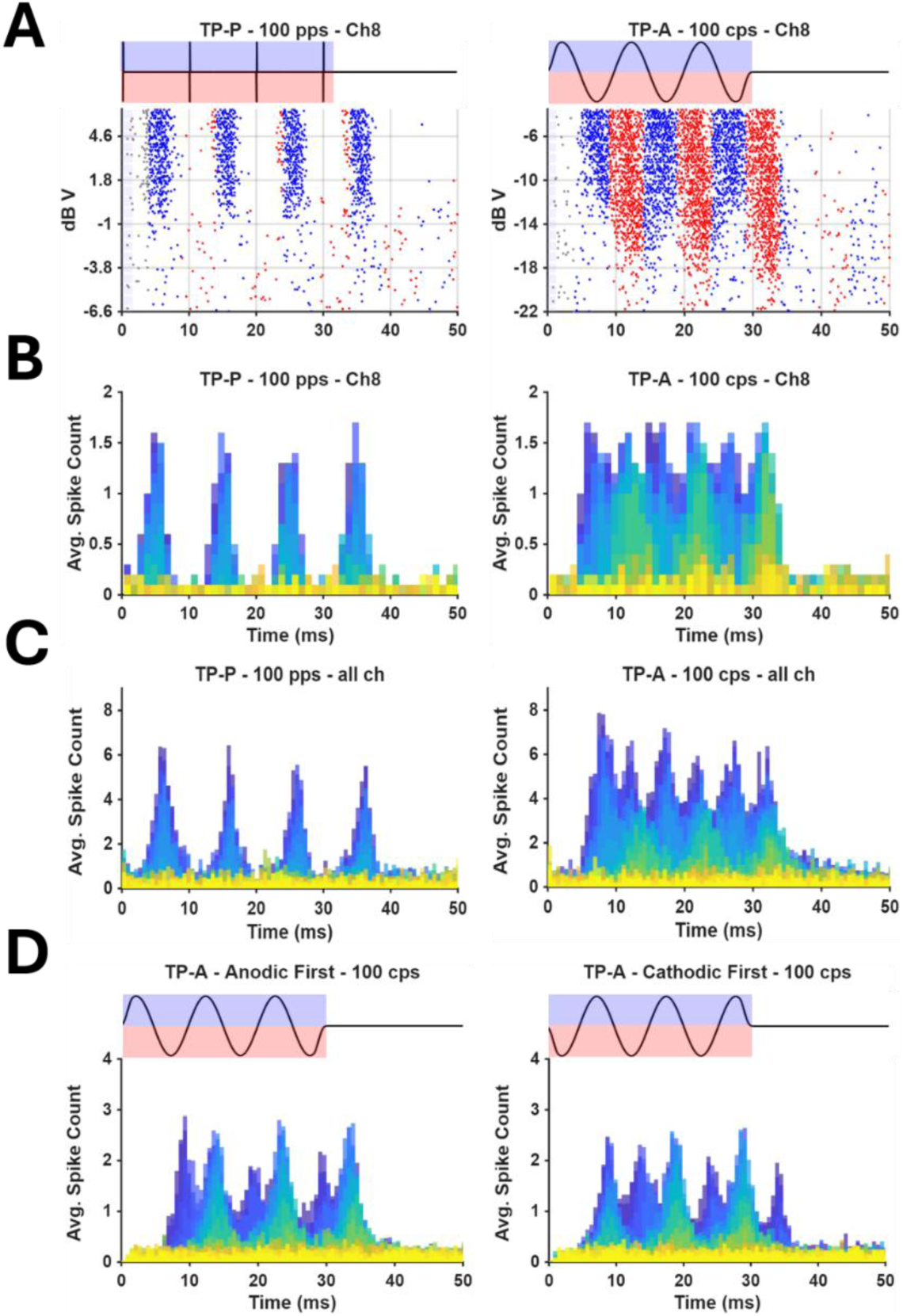
Temporal profile of IC responses. A. Individual recording channel raster plots. At left are responses to a 30 ms tripolar pulse stimulation (TP-P) at 100 pps. At right are same channel’s responses to a 30 ms tripolar analog stimulation (TP-A) at 100 cps. Stimulation waveforms are also reported in black and colored areas delimit phase polarities (blue: anodic; red: cathodic). Alternating colors within rasters temporally represent each phase within stimulation cycles. Grey dots represent events that are most likely not evoked by CI stimulation as their latencies are below 4 ms. B. Individual channel post-stimulus time histograms (PSTH) of responses in A. Color code represents levels of stimulation (Yellow = lowest level of stimulation, dark blue = highest level of stimulation). C. Mean PSTH of all 16 channels in response to stimuli in A. Right panel shows a polarity effect that appeared to be stimulation level dependent. Color code is similar to B. D. Mean PSTH of all 16 channels in response to a 30 ms TP-A stimulation at 100 cps with Anodic-first sinusoids only (left) and with Cathodic-first sinusoids only (right). Color code similar as B and C, stimulation waveforms in black. Note how cathodic phases of the stimulation evoked responses at lower thresholds (green PSTHs) than anodic phases and the appearance of peaks at both cathodic and anodic parts of the stimulation at higher levels of stimulation, confirming a polarity effect in favor of the cathodic phase of the stimulation.

### Analog-evoked responses have higher tonic components but lower synchronization

In general, pulsatile stimulation elicited lower tonic responses than analog stimulation. (Pulsatile: 10 to 15.6% (medians, IQRs [10.2 – 20.52]); Analog: 22.9 to 33% (medians, IQRs [16.89 – 24.6]). An individual example is presented in Fig. 4 where less than 1% of responses are tonic for pulsatile stimulation (Fig. 4.A) but 19.2% are tonic for analog stimulation (Fig. 4.B). The percentage of tonic response was always higher with analog than with pulsatile stimulation for bipolar (p-values < 0.01 for BP-A vs. BP-P) and tripolar (p-values < 0.05 for TP-A vs. TP-P) but not for monopolar (all p-values > 0.05).

**Figure 4:**
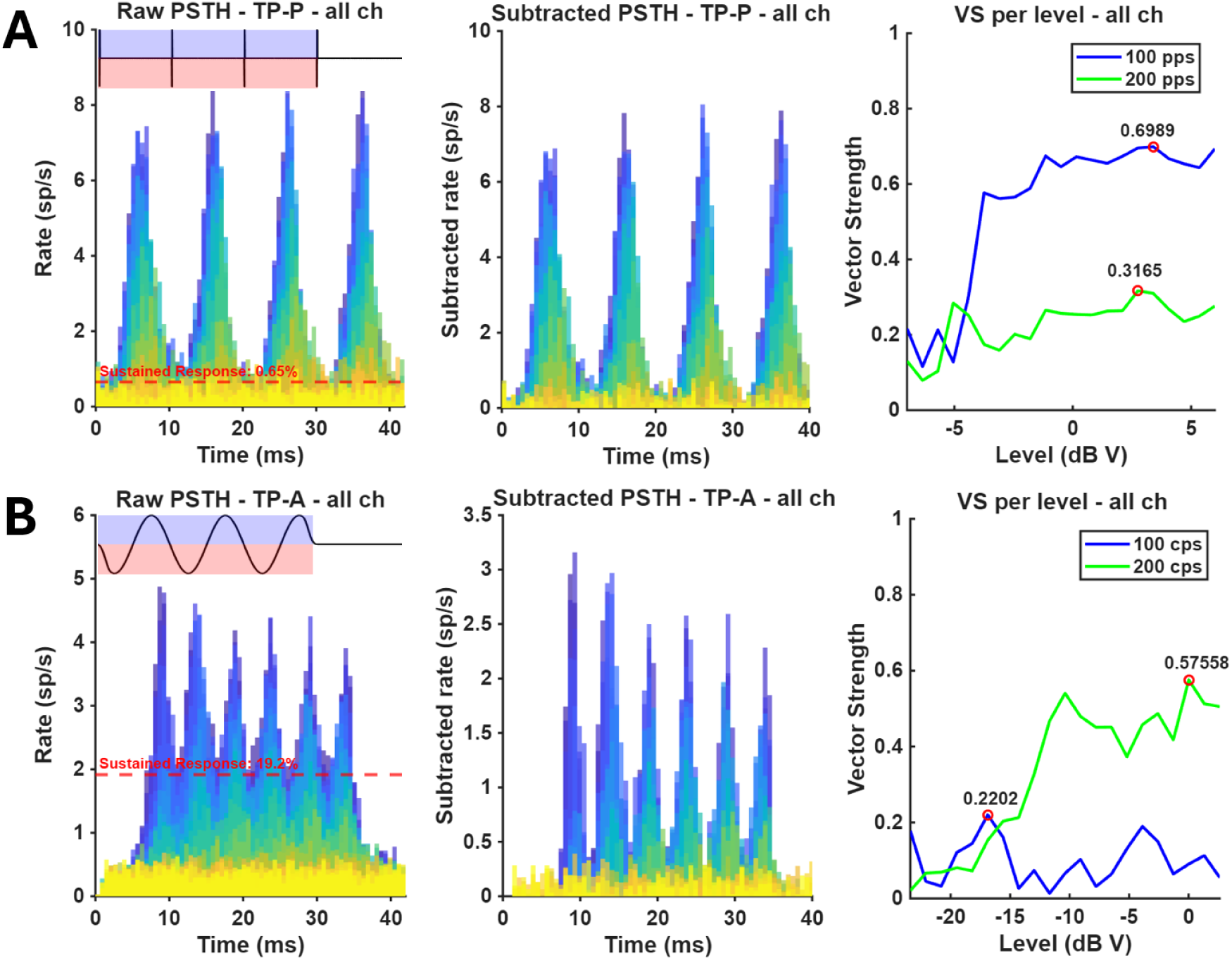
A. Left: Mean post-stimulus time histogram (PSTH) of IC evoked responses by a 30 ms tripolar pulsatile stimulation (TP-P) at 100 pps. Color code represents levels of stimulation from -10 dB V (yellow) to 6 dB V (dark blue) and red dashed line represents the average calculated sustained response (see Methods). Middle: Mean PSTH after subtracting the tonic component of the response. Right: Mean vector strength (VS) of all 16 channels as a function of stimulation level for either 100 pps (blue) or 200 pps (green). The maximum VS (red circle) has been highlighted. B. Left: Mean PSTH of IC evoked responses by a 30 ms tripolar analog stimulation (TP-A) at 100 cps. Color code represents levels of stimulation from -23 dB V (yellow) to 2 dB V (dark blue) and red dashed line represents the average calculated tonic component of the response. Middle: Mean PSTH after subtracting the tonic component of the response. Right: Mean VS of all 16 channels as a function of stimulation level for either 100 cps (Blue) or 200 cps (Green). The maximum VS (red circle) has been highlighted. Note how highest VS is reached for a 200 cps rate at 0.575.

Overall, responses synchrony, as quantified with vector strength (VS) was significantly lower for analog as compared to pulsatile stimulation (Fig. 4, right; Fig. 5). Individual measurements of VS are shown as a function of stimulation levels in Fig. 4, right panels. The VS input/output functions for pulsatile stimulation showed a strong synchrony for 100 pps plateauing around 0.6 to 0.65 with a peak at 0.67. The VS was also calculated for 200 pps for which neural synchrony remained low with increasing stimulus levels (max VS = 0.32). For analog stimulation, the VS I/O functions after subtracting the tonic responses showed a strong synchronization for 200 cps (max VS = 0.58) but not for 100 cps (max VS = 0.22). Figure 5.A also presents individual examples of VS but as a function of stimulation level across the IC recording array. Each panel corresponds to one stimulation paradigm used in this study (Left column: pulsatile, right column: analog; Top: MP, middle: BP and bottom: TP) and color code represents VS values (blue = 0, red = 1). Examples are from one animal and use the same active CI electrode. Group data reports lower VSs for analog when stimulating in bipolar (Fig 5.B, top, BP-P: 0.785 ± 0.132, BP-A: 0.454 ± 0.195, p-value = 0.0044) and tripolar modes (TP-P: 0.744 ± 0.199, TP-A: 0.489 ± 0.203, p-value = 0.0159) but were not significantly different with monopolar mode (MP-P: 0.723 ± 0.216, MP-A: 0.488 ± 0.222, p-value = 0.1289). Synchrony was also equivalent across modes of stimulation with AS (MP-A vs. BP-A vs. TP-A, all p-values < 0.05).

**Figure 5:**
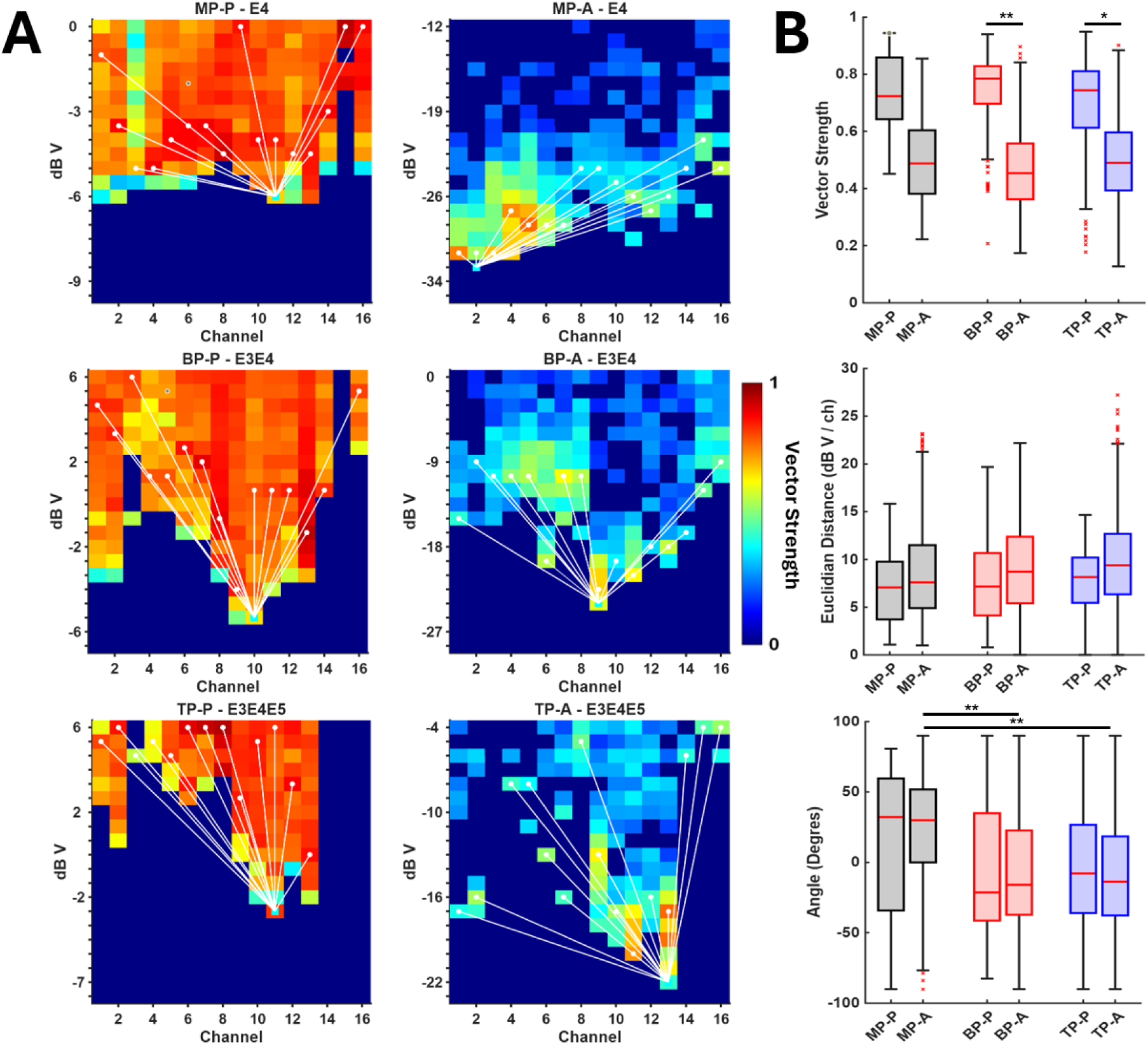
A. Left: Heatmaps representing vector strength as a function of stimulation level for each along the channel array for each pulsatile mode of stimulation used in this study (MP-P, BP-P and TP-P; top, middle and bottom respectively). Red colors represent high values of vector strength and thus, good synchrony for 100 pps stimulation rate. Maximum registered VS under each channel is represented by white dots and Euclidian distances to the threshold by white lines. Right: Similar representation of vector strength but for the associated analog modes of stimulation at a 100 cps (MP-A, BP-A and TP-A; top, middle and bottom respectively). B. Top: Distribution of vector strength observed in the IC across tested modes of stimulation (grey: monopolar, red: bipolar, blue: tripolar; nomenclature is identical to Fig 2). Middle: Distribution of Euclidian distances observed in the IC across tested modes of stimulation. Bottom: Distribution of Euclidian angles observed in the IC across tested modes of stimulation. All statistical tests were hierarchical permutation tests and significance is reported through p-values as follow, *p<0.05, **p<0.01, ***p<0.001.

To further characterize synchrony across the tonotopic gradient, Euclidian Distances (EuD) and Angles (EuA) were calculated across channels within each 16-channel recording (see Methods). In theory, EuDs represented the spatial organization of synchrony along the tonotopic axis while EuAs provided information about the sharpness of VS “tuning” and its directionality. In individual examples provided in Figure 5.A, the maximum VS of each recording channel is designated by a white dot while the lowest response threshold within the array is represented as a blue square. Euclidian distances and Theta angles in respect to this threshold are represented by the white lines connecting white dots to the blue square. The spatial organization of maximum synchrony along the tonotopic axis, represented by the EuDs (Fig 5.B, middle) did not significantly differ between pulsatile and analog stimulation, regardless of the stimulation mode (all p-values < 0.05). Finally, EuAs did not differ significantly between pulsatile and analog within the same mode of stimulation (Fig 5.B, bottom, MP-P: 32.16° ± 93.74, MP-A: 29.98° ± 51.69, p-value = 0.874; BP-P: -21.41° ± 76.39, BP-A: -15.94° ± 59.92, p-value = 0.844; TP-P: -7.91° ± 62.81, TP-A: -13.77° ± 56.18, p-value = 0.54). EuAs were sharper with bipolar and tripolar analog stimulations compared to monopolar analog (MP-A vs. BP-A, MP-A vs. TP-A, p-values = 0.009 and 0.004, respectively), although this result was expected as receptive fields were naturally narrowing due to the more focused stimulation mode used. Thus, although the VSs as a function of stimulation level differed between pulsatile and analog stimulation strategies, the EuDs and EuAs did not capture these differences. One hypothesis would be that it did not account for the nonmonotonicity and/or the polarity effect observed within responses to analog stimulation. After the VS was calculated for both 100 and 200 Hz on all recordings, the ratio VS200 Hz/VS100 Hz was plotted as a function of the maximum VS100 Hz measured within 16-channel array (Fig. 6) for pulsatile (red dots) and analog (blue dots). Data in response to Monopolar, Bipolar and Tripolar stimulation are shown from left to right. Ratios greater than 1 indicate that VS was stronger for 200 Hz than for 100 Hz. For all three stimulus configurations the data with VS ratios greater than 1 are almost all from responses to analog stimulation.

**Figure 6:**
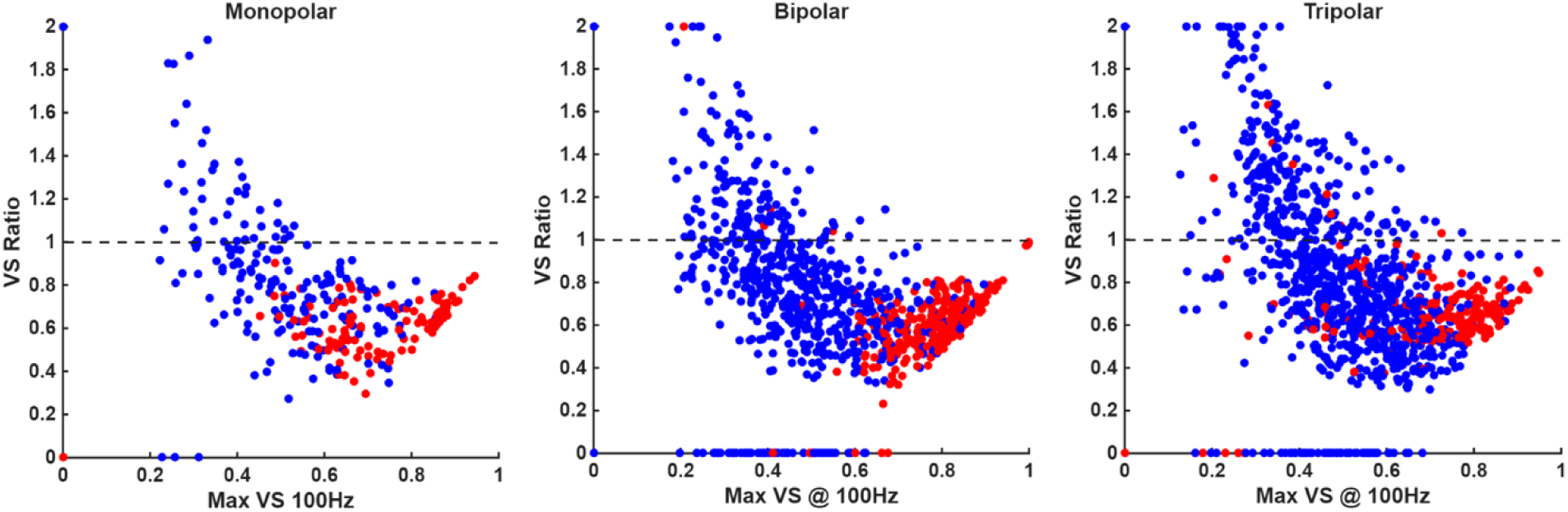
Scatter plots reporting vector strength ratios (VS200 Hz/VS100 Hz) as a function of the maximum measured VS for 100 Hz when stimulating in monopolar (left), bipolar (middle) and tripolar (right) mode. Each dot represents a recording channel with either 100 pps pulsatile-evoked responses (red dots) or 100 cps analog-evoked responses (blue dots). A VS ratio value equal to 1 (marked by the dashed gray line) means that evoked responses had equivalent synchrony for 100 Hz and 200 Hz stimulation rate. Values higher than 1 mean that evoked response had higher synchrony for 200 Hz than for 100 Hz whereas VS Ratio values lower than 1 indicate evoked response more strongly synchronized to 100 Hz than 200 Hz. VS Ratio values were caped to 2 to avoid outliers’ distortion. Note that most dots above 1 are linked to analog stimulation.

## Discussion

The present study evaluated for the first time the coupling of sinusoidal analog stimulation with current focusing modes (bipolar, tripolar) with the objective of comparing IC responses to analog with more conventional pulsatile stimulation. Results replicated previous findings related to electrode configuration, namely that spread of activation narrowed (19,29), and threshold of activation increased systematically with more focused modes of stimulation (from monopolar, to bipolar, to tripolar) (28,42). Within mode of stimulation, analog stimulation always elicited responses with lower thresholds but similar spread of activation, apart from bipolar, which was narrower with analog stimulation. Lastly, IC responses to analog stimulation differed from pulsatile in three ways: 1) lower synchrony, 2) higher sustained/tonic activity, and 3) a level-dependent polarity effect.

### I. Temporal integration governs thresholds

Previous studies have not directly compared responses to continuous sinusoids with pulse trains using neurophysiology experiments in animal models. However, past experiments in cats compared sinusoidally amplitude-modulated biphasic pulse trains stimulation to unmodulated pulse trains (43). Note that stimuli used in that study consisted of longer phase duration pulses of 200 µs compared to the 40 µs used in the present study. These methodological differences make results of this previous study difficult to compare with the present ones.

In the present study, thresholds for pulsatile stimulation were, on average, 17 dB higher than for sinusoidal stimulation. Psychoacoustic studies have demonstrated that detection thresholds decrease systematically with increasing stimulus duration, reflecting the integration of stimulus energy over time (44–46). Heil and Neubauer (47) proposed that auditory detection depends on the accumulation of neural excitation energy within a fixed temporal integration window, typically on the order of tens of milliseconds. Although electric stimulation bypasses cochlear transduction and hair cell-synapses processing, previous psychophysical and physiological studies suggest that temporal integration remains a fundamental constraint in electric hearing (48–52). While analog stimulation delivers energy continuously throughout each cycle, pulsatile stimulation concentrates energy into brief microsecond-scale phases separated by silent intervals, thus requiring substantially higher peak amplitudes to achieve equivalent integrated energy. In the present study, at the level of the IC, this principle is extended and results demonstrate differences quantitatively consistent with predictions from energy-based detection models cited above. This supports the interpretation that system-level temporal integration dominates neural thresholds in the electrically stimulated auditory system.

Despite the pronounced threshold differences, analog and pulsatile stimulation did not significantly differ in their spread of excitation widths within a given mode of stimulation (apart from bipolar). We note, however, that there is a significant narrowing of the spread of excitation for bipolar with analog stimulation and a trend with monopolar. The MP and TP spread of excitations may be at ceiling and floor, respectively, and therefore the experimental paradigm is intrinsically limited in its ability to detect subtle differences in SoE between the stimulation waveforms. The analog stimulation at lower level may be sometimes narrower because of the reduction in thresholds leading to activation from narrower current spread in the cochlea. Alternatively, the relatively small changes in spread of activation may indicate that temporal and spatial aspects of neural activation might be governed by independent processes. Spatial tuning under electrical stimulation is primarily determined by the spatial distribution of the electric field, which depends on electrode geometry, tissue conductivity and stimulation configuration (53–56). Since analog and pulsatile stimulations employed identical electrode configurations and differed only in waveform shape, they most likely generated similar voltage gradients/current spread within the cochlea. Consequently, increasing stimulus amplitude to compensate for reduced temporal integration under pulsatile stimulation primarily scaled neural activation across space, without substantially altering the spatial excitation profile. This dissociation between threshold and spatial tuning has been reported previously in psychophysical studies with CI users, in which increasing the stimulus level of a target pulse train did not increase the breadth of perceptual tuning (57,58). In summary, threshold reduction may not always result in narrower spread of excitation and the physiological data in this study provides direct neural confirmation of this principle at the level of the IC for the MP and TP modes, although this may be because of the limitations of our measures. These findings suggest that substantial waveform-dependent threshold shifts can occur without measurable changes in spread of activation, at least when using the methods employed in the present study for measuring spread of excitation.

### II. Temporal synchrony and phase locking: Pulsatile vs. Analog stimulation

The present study examined pulse trains and analog stimulation with a rate of 100 Hz (or pps/cps), below the 200-300 Hz (pps/cps) upper limit of phase locking to unmodulated pulse trains at the level of the IC where rate coding may be relied on to discriminate rates. The present results of lower vector strength and increased variability in spike timing with analog stimulation are in line with early studies at the level of the auditory nerve fiber suggesting that, unlike square pulses, analog stimulation produces temporal response profiles more similar to those of acoustic pure tones (59–61). Reduced neural synchrony likely occurs through gradual membrane depolarization, allowing intrinsic membrane filtering and stochastic channel fluctuations to introduce temporal dispersion, thereby mitigating the unnatural synchrony inherent to pulsatile cochlear implant stimulation. As reviewed in Carlyon et al., 2025 (27, section “Bertrand Delgutte and Yoojin Chung”), the over-synchrony across auditory nerve (AN) fibers (or the decrease in stochasticity in their responses) evoked by electric stimulation is thought to be the root of distortions in temporal envelope coding, pitch perception, more complex sound appreciation in cochlear implant users, specifically in binaural settings. Briefly, current spread induces the synchronization of large populations of neurons, rendering impossible for AN fibers to follow the “volley principle”, becoming synchronized to each other (63). Kiang and Moxon (64), also observed that electrical stimulation elicits abnormally high neural synchrony compared to acoustic stimulation. Single AN fiber recordings demonstrated phase locking of fibers up to 5 kHz with acoustic pure tone stimulation (65) whereas with electric stimulation (pulse and sinusoid alike) AN fibers could phase-lock above 10 kHz (66–68). Thus, electrical stimulation elicits an unnatural degree of synchronization associated with reduced jitter in the timing of action potentials.

In the present study, we measured multi-unit responses from the IC, as opposed to single nerve fibers in the auditory nerve, and yet we observed responses with a high level of synchrony, especially for unmodulated pulse trains. Pulse train evoked responses were constrained to a small fraction of the stimulus cycle because they were synchronized to the biphasic pulses while analog-evoked responses were distributed across either half or the entire cycle because of the gradual changes in current intensity described above. This led to an increase in sustained or tonic responses with analog stimulation. Synchronous and tonic activities in the IC arise from convergent excitatory inputs and local inhibitory–excitatory interactions that reshape temporal coding (69) but without isolating single units, it remains difficult to prove that the observed tonic component of the MUAs emerged from neurons having a sustained, continuous firing through the cycle or that it originated from several neurons firing at different cycle sections or even different cycles. Nevertheless, tonic firing in response to high stimulus-level electrical stimulation has been documented in both the auditory nerve and IC (64,70). This may reflect sustained depolarization, broadened neural recruitment, and/or integration within midbrain circuitry. Although the emergence of a strong tonic component at higher stimulation levels could be thought of as detrimental to the temporal structure of the response, with sinusoidal amplitude-modulated tones (SAM), a substantial population of IC neurons present a combination of onset and sustained response profiles (71).

One comparison that was not included in the present study is one of SAM pulse trains vs. sinusoidal analog. However, Hancock et al. (2017) (72) reported that neuronal spiking was tightly confined to a narrow temporal window with SAM, specifically mentioning responses to SAM pulse trains to be similar to unmodulated pulse trains, which would be equivalent to the pulse trains used in this study. It is also of note that SAM noises and tones have been described as poor stimuli to reproduce the diversity of envelopes existing in natural acoustic sounds (73), such that SAM-evoked responses in the IC are missing critical non-linearities needed to process complex inputs such as speech or reverberant sounds (74–76). Thus, analog appears to promote more distributed spike timing and features sustained responses profiles that are largely absent from pulsatile stimulation, which may allow for level-dependent encoding of stimuli as well as temporal fine structure. This might improve population-level coding of electrical signals and partially restore natural temporal dispersion.

### III. Polarity-Dependent Responses, Apparent Frequency Doubling, and Implications for Pitch Coding

A mechanistically informative finding of the present study is the strong polarity dependence of IC responses to analog stimulation and its interaction with stimulation level. At low stimulation levels, neural responses were predominantly evoked during the cathodic phase of the waveform, consistent with classical biophysical models showing that cathodic current preferentially depolarizes peripheral processes of AN fibers (55,77,78). At higher stimulation levels, however, we observed robust spiking during both cathodic and anodic phases, leading to neural synchronization at twice the nominal stimulus frequency (i.e., an apparent 200 Hz response to a 100 Hz sinusoidal waveform). This polarity dominance has been repeatedly observed in animal models (79–81). Briefly, it is known that SGNs peripheral processes are more sensitive to cathodic currents whereas their somas are preferentially depolarized by anodic currents. Since peripheral processes usually sit closer to the CI array, cathodic-first pulses are associated with lower thresholds. Those results prompted the idea that the difference in thresholds for cathodic and anodic stimuli, referred to as the polarity effect (PE) could be used to probe the presence or absence of peripheral processes (82). Similarly, polarity-dependent recruitment shifts with increasing levels have been reported in the studies cited above, where higher current amplitudes reduce the asymmetry between anodic and cathodic excitation by recruiting additional neural elements, including central axons and nodes of Ranvier.

Hair cells do not show synaptic activation to both phases of a pure tone, with only one direction of the ciliary deflection being excitatory. Thus, sound-evoked neural responses only arise from one of the two phases of pure tones (83,84). On the other hand, it is well established that electric stimulation, by merit of bypassing hair cells signal transduction, stimulates neurons efficiently with both phases (59,67,85,86). For low-rate sinusoidal stimulation, the inter-peak interval of each phase is long enough to trigger at least one spike per phase and provides neural activation whereas for pulsatile stimulation of similar rate, pulse phases are too close to each other to avoid such dual effects. Psychophysical work indicates that pitch perception under electrical stimulation depends primarily on the fundamental periodicity of the waveform envelope, rather than on the presence of additional phase-locked discharges within each cycle. Stupak et al. 2018 (16) demonstrated with a multidimensional scaling approach that CI users did not report differences in pitch when stimulated with analog compared to pulsatile stimulation at 100 Hz (pps/cps). Their study suggests that, despite neural responses doubling in rate for higher stimulation levels with analog sinusoids, pitch perception remained anchored to 100 Hz This dissociation suggests that central pitch extraction mechanisms likely do not rely purely on the phase-locking of neurons in the IC. Early work in human perception also demonstrated that perceived pitch increases with pulse rate even in the absence of spectral cues (87). Subsequent studies with pulsatile strategies confirmed that CI listeners discriminate pitch primarily based on stimulation rate across a range of conditions (88) and have further shown that complex pulse patterns or waveform manipulations do not substantially alter pitch perception so long as the fundamental temporal periodicity is preserved (89,90). These findings suggest that although analog stimulation evokes neural responses synchronized to both phases of the waveform at high levels, the perceptual pitch would still be expected to correspond to the stimulus frequency. The coexistence of tonic firing and doubled phase locking in a level-dependent manner suggests that analog stimulation engages multiple coding regimes simultaneously, including envelope-based periodicity coding, polarity-driven harmonic components, and sustained rate coding.

### IV. Implications and future directions

The present findings carry important implications for both auditory neuroscience and CI engineering and development. It is worth noting that the translation of focused stimulation to functional performance gains in CI users has been inconsistent. For example, bipolar or tripolar mode with pulsatile stimulation requires higher current to reach threshold and therefore more battery consumption (91). A compounding issue is that focused stimulation is not necessarily beneficial at all electrode locations due to irregular local nerve survival patterns (92,93). The observation that analog stimulation coupled with current focusing configurations, achieved lower thresholds of neural activation suggests that it could improve stimulation efficiency, potentially reduce power consumption, channel interaction, and current spread, while preserving the temporal representation of sound. Current CI systems rely almost exclusively on brief biphasic pulses, largely for safety and technical simplicity. However, this design inherently sacrifices temporal energy integration efficiency and the encoding of temporal fine structure. Our results suggest that alternative stimulation paradigms employing analog waveforms may achieve equivalent neural activation with significantly lower peak currents, consistent with prior modeling and physiological studies (61,70). Furthermore, the invariance of spatial tuning width with tripolar, and possible sharpening with bipolar and monopolar, indicates that such strategies need not compromise spectral resolution, a critical consideration for speech and music perception in CI users. Collectively, these findings reveal potential trade-offs between waveform structure, synchrony, and coding fidelity. Pulsatile stimulation ensures strong, temporally precise neural activation but promotes hypersynchrony. Analog stimulation reduces hypersynchrony and introduces level-dependent sustained dynamics that may more closely approximate acoustic activation patterns, yet it can generate complex nonlinear response transformations at higher levels that are still not fully understood. It is of great interest to revisit analog stimulation in combination with these recent advancements and modern current focusing. It is also worth pursuing more fundamental questions on temporal integration with CIs for conveying temporal fine structure for understanding speech in noise or appreciating music. More natural IC response properties for analog stimulation could also be beneficial for transmitting an important cue for sound localization that is often a challenge for individuals with CIs, that is inter-aural time differences (62).

## Acknowledgments

This work was funded by a grant from the National Institutes of Health R01DC019916 and the generous support of the Ansin Foundation. The authors would like to thank Dr. Ken Hancock, Carter Tims and the Eaton-Peabody Labs engineering core at Mass. Eye and Ear for their assistance and material help; Drs. Shelley Fried, David Lansdberger, Natalia Stupak and Yoojin Chun for their insightful discussions and comments. The authors also dedicate this study to the memory of Pr. Bertrand Delgutte.

## Notes

### Competing Interest Statement

The authors have declared no competing interest.

### Summary of Updates

This version of the MS has updated figures and text to facilitate comprehension. The MS now includes an acknowledgment section. Various typos have been corrected.

## References

1. Wilson BS, Dorman MF. Cochlear implants: a remarkable past and a brilliant future. Hear Res. 2008 Aug;242(0):3–21. doi:10.1016/j.heares.2008.06.005 PubMed PMID: 18616994; PubMed Central PMCID: PMC3707130.

2. Zeng FG. Temporal pitch in electric hearing. Hear Res. 2002 Dec 1;174(1):101–6. doi:10.1016/S0378-5955(02)00644-5

3. Zeng FG, Tang Q, Lu T. Abnormal Pitch Perception Produced by Cochlear Implant Stimulation. PLOS ONE. 2014 Feb 13;9(2):e88662. doi:10.1371/journal.pone.0088662

4. Does cochlear implantation restore music appreciation? - Kohlberg - 2014 - The Laryngoscope - Wiley Online Library [Internet]. [cited 2026 Apr 22]. Available from: https://onlinelibrary.wiley.com/doi/10.1002/lary.24171

5. Bruns L, Mürbe D, Hahne A. Understanding music with cochlear implants. Sci Rep. 2016 Aug 25;6:32026. doi:10.1038/srep32026 PubMed PMID: 27558546; PubMed Central PMCID: PMC4997320.

6. Wilson BS, Finley CC, Lawson DT, Wolford RD, Eddington DK, Rabinowitz WM. Better speech recognition with cochlear implants. Nature. 1991 Jul 18;352(6332):236–8. doi:10.1038/352236a0 PubMed PMID: 1857418.

7. Verschooten E, Shamma S, Oxenham AJ, Moore BCJ, Joris PX, Heinz MG, et al. The upper frequency limit for the use of phase locking to code temporal fine structure in humans: A compilation of viewpoints. Hear Res. 2019 Jun 1;377:109–21. doi:10.1016/j.heares.2019.03.011 PubMed PMID: 30927686; PubMed Central PMCID: PMC6524635.

8. Riss D, Hamzavi JS, Blineder M, Honeder C, Ehrenreich I, Kaider A, et al. FS4, FS4-p, and FSP: a 4-month crossover study of 3 fine structure sound-coding strategies. Ear Hear. 2014;35(6):e272–281. doi:10.1097/AUD.0000000000000063 PubMed PMID: 25127325.

9. Fischer T, Schmid C, Kompis M, Mantokoudis G, Caversaccio M, Wimmer W. Effects of temporal fine structure preservation on spatial hearing in bilateral cochlear implant users. J Acoust Soc Am. 2021 Aug 1;150(2):673–86. doi:10.1121/10.0005732 PubMed PMID: 34470279.

10. Shannon RV, Zeng FG, Kamath V, Wygonski J, Et. A. Speech Recognition with Primarily Temporal Cues. Science. 1995;270(5234):5234. doi:10.1126/science.270.5234.303 PubMed PMID: 7569981.

11. Landsberger DM, McKay CM. Perceptual differences between low and high rates of stimulation on single electrodes for cochlear implantees. J Acoust Soc Am. 2005 Jan;117(1):319– 27. doi:10.1121/1.1830672 PubMed PMID: 15704424.

12. Goldsworthy RL, Shannon RV. Training improves cochlear implant rate discrimination on a psychophysical task. J Acoust Soc Am. 2014 Jan 1;135(1):334–41. doi:10.1121/1.4835735

13. Hochmair-Desoyer IJ, Hochmair ES, Burian K, Stiglbrunner HK. Percepts from the Vienna cochlear prosthesis. Ann N Y Acad Sci. 1983;405:295–306. doi:10.1111/j.1749-6632.1983.tb31642.x PubMed PMID: 6223552.

14. Hochmair ES, Hochmair-Desoyer IJ, Stiglbrunner HK. Psychoacoustic temporal speech understanding in cochlear implant patients. In: M. M. SRA et M, editor. Cochlear Implants. New York: Raven Press; 1984. p. 291–304.

15. House WF, Urban J. Long term results of electrode implantation and electronic stimulation of the cochlea in man. Ann Otol Rhinol Laryngol. 1973;82(4):504–17. doi:10.1177/000348947308200408 PubMed PMID: 4721186.

16. Stupak N, Padilla M, Morse RP, Landsberger DM. Perceptual Differences Between Low-Frequency Analog and Pulsatile Stimulation as Shown by Single- and Multidimensional Scaling. Trends Hear. 2018;22:2331216518807535. doi:10.1177/2331216518807535 PubMed PMID: 30378468; PubMed Central PMCID: PMC6236864.

17. Koch DB, Osberger MJ, Segel P, Kessler D. HiResolution and conventional sound processing in the HiResolution bionic ear: using appropriate outcome measures to assess speech recognition ability. Audiol Neurootol. 2004;9(4):214–23. doi:10.1159/000078391 PubMed PMID: 15205549.

18. Battmer RD, Zilberman Y, Haake P, Lenarz T. Simultaneous Analog Stimulation (SAS)--Continuous Interleaved Sampler (CIS) pilot comparison study in Europe. Ann Otol Rhinol Laryngol Suppl. 1999 Apr;177:69–73. doi:10.1177/00034894991080s414 PubMed PMID: 10214805.

19. Snyder RL, Bierer JA, Middlebrooks JC. Topographic Spread of Inferior Colliculus Activation in Response to Acoustic and Intracochlear Electric Stimulation. JARO J Assoc Res Otolaryngol. 2004 Sep;5(3):3. doi:10.1007/s10162-004-4026-5 PubMed PMID: 15492888; PubMed Central PMCID: PMC2504547.

20. Srinivasan AG, Padilla M, Shannon RV, Landsberger DM. Improving speech perception in noise with current focusing in cochlear implant users. Hear Res. 2013 mai;299:29–36. doi:10.1016/j.heares.2013.02.004

21. Bierer JA, Middlebrooks JC. Cortical responses to cochlear implant stimulation: channel interactions. J Assoc Res Otolaryngol JARO. 2004 Mar;5(1):32–48. doi:10.1007/s10162-003-3057-7 PubMed PMID: 14564662; PubMed Central PMCID: PMC2538368.

22. Quass GL, Kral A. Tripolar configuration and pulse shape in cochlear implants reduce channel interactions in the temporal domain. Hear Res. 2024 Mar 1;443:108953. doi:10.1016/j.heares.2024.108953 PubMed PMID: 38277881.

23. Imennov NS, Won JH, Drennan WR, Jameyson E, Rubinstein JT. Detection of acoustic temporal fine structure by cochlear implant listeners: behavioral results and computational modeling. Hear Res. 2013 Apr;298:60–72. doi:10.1016/j.heares.2013.01.004 PubMed PMID: 23333260; PubMed Central PMCID: PMC3605703.

24. Gourévitch B, Edeline JM. Age-related changes in the guinea pig auditory cortex: relationship with brainstem changes and comparison with tone-induced hearing loss. Eur J Neurosci. 2011 Dec;34(12):12. doi:10.1111/j.1460-9568.2011.07905.x PubMed PMID: 22092590.

25. Aushana Y, Souffi S, Edeline JM, Lorenzi C, Huetz C. Robust Neuronal Discrimination in Primary Auditory Cortex Despite Degradations of Spectro-temporal Acoustic Details: Comparison Between Guinea Pigs with Normal Hearing and Mild Age-Related Hearing Loss. J Assoc Res Otolaryngol JARO. 2018 Apr;19(2):163–80. doi:10.1007/s10162-017-0649-1 PubMed PMID: 29302822; PubMed Central PMCID: PMC5878150.

26. Sato M, Baumhoff P, Kral A. Cochlear Implant Stimulation of a Hearing Ear Generates Separate Electrophonic and Electroneural Responses. J Neurosci Off J Soc Neurosci. 2016 Jan 6;36(1):1. doi:10.1523/JNEUROSCI.2968-15.2016 PubMed PMID: 26740649.

27. Sato M, Baumhoff P, Tillein J, Kral A. Physiological Mechanisms in Combined Electric–Acoustic Stimulation. Otol Neurotol. 2017 Sep;38(8):e215–23. doi:10.1097/MAO.0000000000001428

28. Kral A, Hartmann R, Mortazavi D, Klinke R. Spatial resolution of cochlear implants: the electrical field and excitation of auditory afferents. Hear Res. 1998 Jul 1;121(1):1–2. doi:10.1016/S0378-5955(98)00061-6 PubMed PMID: 9682804.

29. Snyder RL, Middlebrooks JC, Bonham BH. COCHLEAR IMPLANT ELECTRODE CONFIGURATION EFFECTS ON ACTIVATION THRESHOLD AND TONOTOPIC SELECTIVITY. Hear Res. 2008 Jan;235(1–2):1–2. doi:10.1016/j.heares.2007.09.013 PubMed PMID: 18037252; PubMed Central PMCID: PMC2387102.

30. McInturff S, Coen FV, Hight AE, Tarabichi O, Kanumuri VV, Vachicouras N, et al. Comparison of Responses to DCN vs. VCN Stimulation in a Mouse Model of the Auditory Brainstem Implant (ABI). J Assoc Res Otolaryngol JARO. 2022 Jun;23(3):391–412. doi:10.1007/s10162-022-00840-8 PubMed PMID: 35381872; PubMed Central PMCID: PMC9085982.

31. McInturff S, Adenis V, Coen FV, Lacour SP, Lee DJ, Brown MC. Sensitivity to Pulse Rate and Amplitude Modulation in an Animal Model of the Auditory Brainstem Implant (ABI). J Assoc Res Otolaryngol. 2023 May 8;24(3):365–84. doi:10.1007/s10162-023-00897-z

32. Dieter A, Duque-Afonso CJ, Rankovic V, Jeschke M, Moser T. Near physiological spectral selectivity of cochlear optogenetics. Nat Commun. 2019 Apr 29;10(1):1962. doi:10.1038/s41467-019-09980-7 PubMed PMID: 31036812; PubMed Central PMCID: PMC6488702.

33. Rees A, Palmer AR. Neuronal responses to amplitude-modulated and pure-tone stimuli in the guinea pig inferior colliculus, and their modification by broadband noise. J Acoust Soc Am. 1989 May 1;85(5):5. doi:10.1121/1.397851

34. Garcia-Lazaro JA, Belliveau LAC, Lesica NA. Independent Population Coding of Speech with Sub-Millisecond Precision. J Neurosci. 2013 Dec 4;33(49):19362–72. doi:10.1523/JNEUROSCI.3711-13.2013 PubMed PMID: 24305831.

35. Bierer JA, Bierer SM, Middlebrooks JC. Partial tripolar cochlear implant stimulation: Spread of excitation and forward masking in the inferior colliculus. Hear Res. 2010 Dec 1;270(1–2):1–2. doi:10.1016/j.heares.2010.08.006 PubMed PMID: 20727397; PubMed Central PMCID: PMC2997905.

36. Stanislaw H, Todorov N. Calculation of signal detection theory measures. Behav Res Methods Instrum Comput J Psychon Soc Inc. 1999 Feb;31(1):137–49. doi:10.3758/bf03207704 PubMed PMID: 10495845.

37. Goldberg JM, Brown PB. Response of binaural neurons of dog superior olivary complex to dichotic tonal stimuli: some physiological mechanisms of sound localization. J Neurophysiol. 1969 Jul;32(4):4. doi:10.1152/jn.1969.32.4.613 PubMed PMID: 5810617.

38. Yin TC, Kuwada S. Binaural interaction in low-frequency neurons in inferior colliculus of the cat. II. Effects of changing rate and direction of interaural phase. J Neurophysiol. 1983 Oct;50(4):1000–19. doi:10.1152/jn.1983.50.4.1000 PubMed PMID: 6631458.

39. Yin TC, Kuwada S. Binaural interaction in low-frequency neurons in inferior colliculus of the cat. III. Effects of changing frequency. J Neurophysiol. 1983 Oct;50(4):1020–42. doi:10.1152/jn.1983.50.4.1020 PubMed PMID: 6631459.

40. Kuwada S, Yin TC. Binaural interaction in low-frequency neurons in inferior colliculus of the cat. I. Effects of long interaural delays, intensity, and repetition rate on interaural delay function. J Neurophysiol. 1983 Oct;50(4):981–99. doi:10.1152/jn.1983.50.4.981 PubMed PMID: 6631473.

41. Chung Y, Hancock KE, Nam SI, Delgutte B. Coding of Electric Pulse Trains Presented through Cochlear Implants in the Auditory Midbrain of Awake Rabbit: Comparison with Anesthetized Preparations. J Neurosci. 2014 Jan 1;34(1):218–31. doi:10.1523/JNEUROSCI.2084-13.2014 PubMed PMID: 24381283; PubMed Central PMCID: PMC3866485.

42. Bierer JA, Middlebrooks JC. Auditory Cortical Images of Cochlear-Implant Stimuli: Dependence on Electrode Configuration. J Neurophysiol. 2002 Jan 1;87(1):1. PubMed PMID: 11784764.

43. Moore CM, Vollmer M, Leake PA, Snyder RL, Rebscher SJ. The effects of chronic intracochlear electrical stimulation on inferior colliculus spatial representation in adult deafened cats. Hear Res. 2002 Feb;164(1–2):1–2. doi:10.1016/s0378-5955(01)00415-4 PubMed PMID: 11950528.

44. Oxenham AJ. Forward masking: adaptation or integration? J Acoust Soc Am. 2001 Feb;109(2):732–41. doi:10.1121/1.1336501 PubMed PMID: 11248977.

45. Plomp R, Bouman MA. Relation between Hearing Threshold and Duration for Tone Pulses. J Acoust Soc Am. 1959 Jun 1;31(6):749–58. doi:10.1121/1.1907781

46. Viemeister NF, Wakefield GH. Temporal integration and multiple looks. J Acoust Soc Am. 1991 Aug 1;90(2):858–65. doi:10.1121/1.401953

47. Heil P, Neubauer H. A unifying basis of auditory thresholds based on temporal summation. Proc Natl Acad Sci U S A. 2003 May 13;100(10):6151–6. doi:10.1073/pnas.1030017100 PubMed PMID: 12724527; PubMed Central PMCID: PMC156341.

48. Carlyon RP, van Wieringen A, Deeks JM, Long CJ, Lyzenga J, Wouters J. Effect of inter-phase gap on the sensitivity of cochlear implant users to electrical stimulation. Hear Res. 2005 Jul 1;205(1):210–24. doi:10.1016/j.heares.2005.03.021

49. McKay CM, McDermott HJ. The perceptual effects of current pulse duration in electrical stimulation of the auditory nerve. J Acoust Soc Am. 1999 Aug;106(2):2. PubMed PMID: 10462805.

50. McKay CM, McDermott HJ. Loudness perception with pulsatile electrical stimulation: The effect of interpulse intervals. J Acoust Soc Am. 1998 Aug 1;104(2):1061–74. doi:10.1121/1.423316

51. Shannon RV. Threshold functions for electrical stimulation of the human cochlear nucleus. Hear Res. 1989 Jun 15;40(1):173–7. doi:10.1016/0378-5955(89)90110-X

52. Shannon RV. A model of threshold for pulsatile electrical stimulation of cochlear implants. Hear Res. 1989 Jul 1;40(3):197–204. doi:10.1016/0378-5955(89)90160-3

53. Frijns JH, de Snoo SL, ten Kate JH. Spatial selectivity in a rotationally symmetric model of the electrically stimulated cochlea. Hear Res. 1996 May;95(1–2):33–48. doi:10.1016/0378-5955(96)00004-4 PubMed PMID: 8793506.

54. Micco AG, Richter CP. Tissue resistivities determine the current flow in the cochlea. Curr Opin Otolaryngol Head Neck Surg. 2006 Oct;14(5):5. doi:10.1097/01.moo.0000244195.04926.a0 PubMed PMID: 16974151.

55. Rattay F, Lutter P, Felix H. A model of the electrically excited human cochlear neuron. I. Contribution of neural substructures to the generation and propagation of spikes. Hear Res. 2001 Mar;153(1–2):43–63. doi:10.1016/s0378-5955(00)00256-2 PubMed PMID: 11223296.

56. Rattay F, Leao RN, Felix H. A model of the electrically excited human cochlear neuron. II. Influence of the three-dimensional cochlear structure on neural excitability. Hear Res. 2001 Mar;153(1–2):64–79. doi:10.1016/s0378-5955(00)00257-4 PubMed PMID: 11223297.

57. Kreft HA, Donaldson GS, Nelson DA. Effects of pulse rate and electrode array design on intensity discrimination in cochlear implant users. J Acoust Soc Am. 2004 Oct;116(4 Pt 1):4 Pt 1. PubMed PMID: 15532657.

58. Bierer JA, Faulkner KF. Identifying cochlear implant channels with poor electrode-neuron interface: partial tripolar, single-channel thresholds and psychophysical tuning curves. Ear Hear. 2010 Apr;31(2):247–58. doi:10.1097/AUD.0b013e3181c7daf4 PubMed PMID: 20090533; PubMed Central PMCID: PMC2836401.

59. van den Honert C, Stypulkowski PH. Temporal response patterns of single auditory nerve fibers elicited by periodic electrical stimuli. Hear Res. 1987;29(2–3):207–22. doi:10.1016/0378-5955(87)90168-7 PubMed PMID: 3624084.

60. Litvak L, Delgutte B, Eddington D. Auditory nerve fiber responses to electric stimulation: modulated and unmodulated pulse trains. J Acoust Soc Am. 2001 Jul;110(1):368–79. doi:10.1121/1.1375140 PubMed PMID: 11508961; PubMed Central PMCID: PMC2275322.

61. Rubinstein JT, Wilson BS, Finley CC, Abbas PJ. Pseudospontaneous activity: stochastic independence of auditory nerve fibers with electrical stimulation. Hear Res. 1999 Jan 1;127(1):108–18. doi:10.1016/S0378-5955(98)00185-3

62. Carlyon RP, Deeks JM, Delgutte B, Chung Y, Vollmer M, Ohl FW, et al. Limitations on Temporal Processing by Cochlear Implant Users: A Compilation of Viewpoints. Trends Hear. 2025;29:23312165251317006. doi:10.1177/23312165251317006 PubMed PMID: 40095543; PubMed Central PMCID: PMC12076235.

63. Schnupp J, Nelken I, King A. Auditory neuroscience: making sense of sound. Cambridge, Mass.; London: MIT Press; 2012.

64. Kiang NS, Moxon E. PHYSIOLOGICAL CONSIDERATIONS IN ARTIFICIAL STIhIULATION OF THE INNER EAR.

65. Palmer AR, Russell IJ. Phase-locking in the cochlear nerve of the guinea-pig and its relation to the receptor potential of inner hair-cells. Hear Res. 1986;24(1):1–15. doi:10.1016/0378-5955(86)90002-x PubMed PMID: 3759671.

66. Delgutte. Physiological mechanisms of psychophysical masking: Observations from auditory-nerve fibers. J Acoust Soc Am. 1990;87(2):791–809. doi:10.1121/1.398891 PubMed PMID: 2307776.

67. Dynes SB, Delgutte B. Phase-locking of auditory-nerve discharges to sinusoidal electric stimulation of the cochlea. Hear Res. 1992 Feb;58(1):79–90. doi:10.1016/0378-5955(92)90011-b PubMed PMID: 1559909.

68. Miller CA, Hu N, Zhang F, Robinson BK, Abbas PJ. Changes Across Time in the Temporal Responses of Auditory Nerve Fibers Stimulated by Electric Pulse Trains. JARO J Assoc Res Otolaryngol. 2008 Mar;9(1):122–37. doi:10.1007/s10162-007-0108-5 PubMed PMID: 18204987; PubMed Central PMCID: PMC2536806.

69. Krishna BS, Semple MN. Auditory temporal processing: responses to sinusoidally amplitude-modulated tones in the inferior colliculus. J Neurophysiol. 2000 Jul;84(1):1. doi:10.1152/jn.2000.84.1.255 PubMed PMID: 10899201.

70. Middlebrooks JC, Snyder RL. Selective Electrical Stimulation of the Auditory Nerve Activates a Pathway Specialized for High Temporal Acuity. J Neurosci. 2010 Feb 3;30(5):1937–46. doi:10.1523/JNEUROSCI.4949-09.2010 PubMed PMID: 20130202; PubMed Central PMCID: PMC2828779.

71. Nelson PC, Carney LH. Neural rate and timing cues for detection and discrimination of amplitude-modulated tones in the awake rabbit inferior colliculus. J Neurophysiol. 2007 Jan;97(1):522–39. doi:10.1152/jn.00776.2006 PubMed PMID: 17079342; PubMed Central PMCID: PMC2577033.

72. Hancock KE, Chung Y, McKinney MF, Delgutte B. Temporal Envelope Coding by Inferior Colliculus Neurons with Cochlear Implant Stimulation. J Assoc Res Otolaryngol JARO. 2017 Dec;18(6):771–88. doi:10.1007/s10162-017-0638-4 PubMed PMID: 28717877; PubMed Central PMCID: PMC5688046.

73. Singh NC, Theunissen FE. Modulation spectra of natural sounds and ethological theories of auditory processing. J Acoust Soc Am. 2003;114(6):6.

74. Moller AR, Rees A. Dynamic properties of the responses of single neurons in the inferior colliculus of the rat. Hear Res. 1986;24(3):3.

75. Delgutte B, Hammond BM, Cariani PA. Neural coding of the temporal envelope of speech: Relation to modulation transfer functions. In: A. R. Palmer AR, Meddis R, editors. Psychophysical and Physiological Advances in Hearing. London; 1998. p. 595–603.

76. Slama MCC, Delgutte B. Neural Coding of Sound Envelope in Reverberant Environments. J Neurosci. 2015 Mar 11;35(10):4452–68. doi:10.1523/JNEUROSCI.3615-14.2015

77. Ranck JB. Which elements are excited in electrical stimulation of mammalian central nervous system: a review. Brain Res. 1975 Nov 21;98(3):417–40. doi:10.1016/0006-8993(75)90364-9 PubMed PMID: 1102064.

78. Briaire JJ, Frijns JH. Field patterns in a 3D tapered spiral model of the electrically stimulated cochlea. Hear Res. 2000 Oct;148(1–2):1–2. PubMed PMID: 10978822.

79. Shepherd RK, Javel E. Electrical stimulation of the auditory nerve. I. Correlation of physiological responses with cochlear status. Hear Res. 1997 Jun;108(1–2):112–44. doi:10.1016/S0378-5955(97)00046-4

80. Miller AL, Smith DW, Pfingst BE. Across-species comparisons of psychophysical detection thresholds for electrical stimulation of the cochlea: I. Sinusoidal stimuli. Hear Res. 1999 Aug 1;134(1):89–104. doi:10.1016/S0378-5955(99)00072-6

81. Miller AL, Smith DW, Pfingst BE. Across-species comparisons of psychophysical detection thresholds for electrical stimulation of the cochlea: II. Strength-duration functions for single, biphasic pulses. Hear Res. 1999 Sep 1;135(1):1–2. doi:10.1016/S0378-5955(99)00089-1 PubMed PMID: 10491953.

82. Konerding W, Arenberg J, Kral A, Baumhoff P. Cochlear health alters the polarity effect and spike-initiation sites in guinea pigs. Hear Res. 2025 Sep 1;465:109341. doi:10.1016/j.heares.2025.109341 PubMed PMID: 40609481.

83. Hudspeth AJ, Corey DP. Sensitivity, polarity, and conductance change in the response of vertebrate hair cells to controlled mechanical stimuli. Proc Natl Acad Sci U S A. 1977 Jun;74(6):2407–11. doi:10.1073/pnas.74.6.2407 PubMed PMID: 329282; PubMed Central PMCID: PMC432181.

84. Guinan JJ, Salt A, Cheatham MA. Progress in Cochlear Physiology after Békésy. Hear Res. 2012 Nov;293(1–2):12–20. doi:10.1016/j.heares.2012.05.005 PubMed PMID: 22633944; PubMed Central PMCID: PMC3530189.

85. Macherey O, Cazals Y. Effects of Pulse Shape and Polarity on Sensitivity to Cochlear Implant Stimulation: A Chronic Study in Guinea Pigs. Adv Exp Med Biol. 2016;894:133–42. doi:10.1007/978-3-319-25474-6_15 PubMed PMID: 27080654.

86. Hartmann R, Topp G, Klinke R. Discharge patterns of cat primary auditory fibers with electrical stimulation of the cochlea. Hear Res. 1984 Jan 1;13(1):1. doi:10.1016/0378-5955(84)90094-7 PubMed PMID: 6546751.

87. Shannon RV. Multichannel electrical stimulation of the auditory nerve in man. I. Basic psychophysics. Hear Res. 1983 Aug 1;11(2):157–89. doi:10.1016/0378-5955(83)90077-1

88. Carlyon RP, Deeks JM. Limitations on rate discrimination. J Acoust Soc Am. 2002 Sep;112(3 Pt 1):1009–25. doi:10.1121/1.1496766 PubMed PMID: 12243150.

89. Carlyon RP, Deeks JM, McKay CM. Effect of Pulse Rate and Polarity on the Sensitivity of Auditory Brainstem and Cochlear Implant Users to Electrical Stimulation. J Assoc Res Otolaryngol JARO. 2015 Oct;16(5):5. doi:10.1007/s10162-015-0530-z PubMed PMID: 26138501; PubMed Central PMCID: PMC4569604.

90. Macherey O, Deeks JM, Carlyon RP. Extending the limits of place and temporal pitch perception in cochlear implant users. J Assoc Res Otolaryngol JARO. 2011 Apr;12(2):2. doi:10.1007/s10162-010-0248-x PubMed PMID: 21116672; PubMed Central PMCID: PMC3046333.

91. Padilla M, Landsberger DM. Reduction in Spread of Excitation from Current Focusing at Multiple Cochlear Locations in Cochlear Implant Users. Hear Res. 2016 Mar;333:98–107. doi:10.1016/j.heares.2016.01.002 PubMed PMID: 26778546; PubMed Central PMCID: PMC4907334.

92. Zhu Z, Tang Q, Zeng FG, Guan T, Ye D. Cochlear Implant Spatial Selectivity with Monopolar, Bipolar and Tripolar Stimulation. Hear Res. 2012 Jan;283(1–2):1–2. doi:10.1016/j.heares.2011.11.005 PubMed PMID: 22138630; PubMed Central PMCID: PMC3277661.

93. Jahn KN, Arenberg JG. Identifying Cochlear Implant Channels With Relatively Poor Electrode-Neuron Interfaces Using the Electrically Evoked Compound Action Potential. Ear Hear. 2020;41(4):961–73. doi:10.1097/AUD.0000000000000844 PubMed PMID: 31972772; PubMed Central PMCID: PMC10443089.

